# Mechanism of Ca^2+^ transport by ferroportin

**DOI:** 10.1101/2022.08.23.504983

**Authors:** Jiemin Shen, Azaan Saalim Wilbon, Ming Zhou, Yaping Pan

**Affiliations:** Verna and Marrs McLean Department of Biochemistry and Molecular Biology, Baylor College of Medicine, Houston, TX 77030, USA

## Abstract

Ferroportin (Fpn) is a transporter that releases Fe^2+^ from cells and is important for iron homeostasis in circulation. Export of Fe^2+^ by Fpn is coupled to import of H^+^ to maintain charge balance. Although Ca^2+^ was shown to modulate Fe^2+^ transport in Fpn, transport of Ca^2+^ by Fpn has not been demonstrated. Here we show that human Fpn (HsFpn) mediates Ca^2+^ transport, and that the Ca^2+^ transport does not rely on the transport of other ions. We determine the structure of Ca^2+^-bound HsFpn and identify a single Ca^2+^ binding site distinct from the Fe^2+^ binding sites. Further studies validate the Ca^2+^ binding site and show that Ca^2+^ transport is inhibited in the presence of Fe^2+^ but not *vice versa*. Function of Fpn as a Ca^2+^ uniporter in the absence of Fe^2+^ provides a molecular basis for regulations of iron homeostasis by Ca^2+^.

## Introduction

Ferroportin (Fpn), encoded by *SLC40A1* gene, is the only known Fe^2+^ exporter in human (Nemeth and Ganz 2021; Abboud and Haile 2000; Donovan et al. 2000; McKie et al. 2000). Fpn is highly expressed on enterocytes, hepatocytes, and macrophages, and mediates absorption of dietary iron and iron recycled from senescent red blood cells (Drakesmith, Nemeth, and Ganz 2015; Mackenzie and Garrick 2005; Knutson et al. 2005). Activity of Fpn is regulated by hepcidin, a small peptide hormone secreted by hepatocytes, which binds to Fpn and reduces iron export by inhibiting its transport activity and triggering endocytosis of Fpn (Aschemeyer et al. 2018; Nemeth et al. 2004; Nemeth and Ganz 2021). Mutations in Fpn or perturbations to the production of hepcidin cause hereditary hemochromatosis and iron-loading anemias in human (Pietrangelo 2017; Ganz 2013; Ginzburg 2019; Vlasveld et al. 2019).

Fpn is a Fe^2+^/2H^+^ exchanger, i.e., export of one Fe^2+^ is accompanied by import of two H^+^ (Pan et al. 2020). The electroneutral transport mechanism is likely an adaptation to overcome the negative membrane potential that would have significantly hindered export of cations. Structures of mammalian Fpns show that there are two ion binding sites (Billesbolle et al. 2020; Pan et al. 2020), termed Site 1 (S1) and Site 2 (S2), that mediate Fe^2+^ and H^+^ transport (Pan et al. 2020). S1 consists of Asp39 and His43, and S2, Cys326 and His507 (Billesbolle et al. 2020; Pan et al. 2020). Mutational studies showed that while both S1 and S2 are important for H^+^ transport, S2 has a more prominent role in mediating Fe^2+^ transport than S1 (Pan et al. 2020).

Ca^2+^ is known to bind mammalian Fpn (Deshpande et al. 2018) but Ca^2+^ transport by Fpn has not been demonstrated. The role of Ca^2+^ in Fe^2+^ transport remains ambivalent (Li, Yang, and Li 2020; Pan et al. 2020; Billesbolle et al. 2020). In an early study using Fpn expressed in *Xenopus* oocytes, Ca^2+^ was shown to be required for Fe^2+^ transport but was not transported by Fpn (Deshpande et al. 2018). More recently, several studies using purified Fpn reconstituted into liposomes showed Fe^2+^ transport occurs in the absence of Ca^2+^ (Li, Yang, and Li 2020; Pan et al. 2020; Billesbolle et al. 2020), and that Ca^2+^ could potentiate Fe^2+^ transport under certain conditions (Li, Yang, and Li 2020; Billesbolle et al. 2020). In the current study, we focus our efforts on understanding Ca^2+^ transport in Fpn.

## Results

### HsFpn is a Ca^2+^ uniporter

HsFpn was expressed and purified (**Figure 1—figure supplement 1a**) and reconstituted into liposomes for transport assays (**Methods**). Significant Ca^2+^ uptake was observed in proteoliposomes containing HsFpn, as indicated by increased fluorescence of Fluo-4 trapped inside (**Figure 1a** and **Figure 1—figure supplement 1b**). In contrast, control liposomes that do not have HsFpn show minimal change in fluorescence (**Figure 1a**). Since a previous study reported that Fpn does not transport Ca^2+^ (Deshpande et al. 2018), we further examined Ca^2+^ transport in Fpn. We found that Ca^2+^ influx is inhibited by hepcidin and by the fragment of antigen-binding (Fab) of a monoclonal antibody (11F9) (Pan et al. 2020) to HsFpn (manuscript submitted together, **Figure 1a–b**), providing further support that the Ca^2+^ influx is mediated by HsFpn. Specific Ca^2+^ uptake is also observed in human embryonic kidney (HEK) cells expressing HsFpn (**Figure 1c–d**).

**Figure 1.**
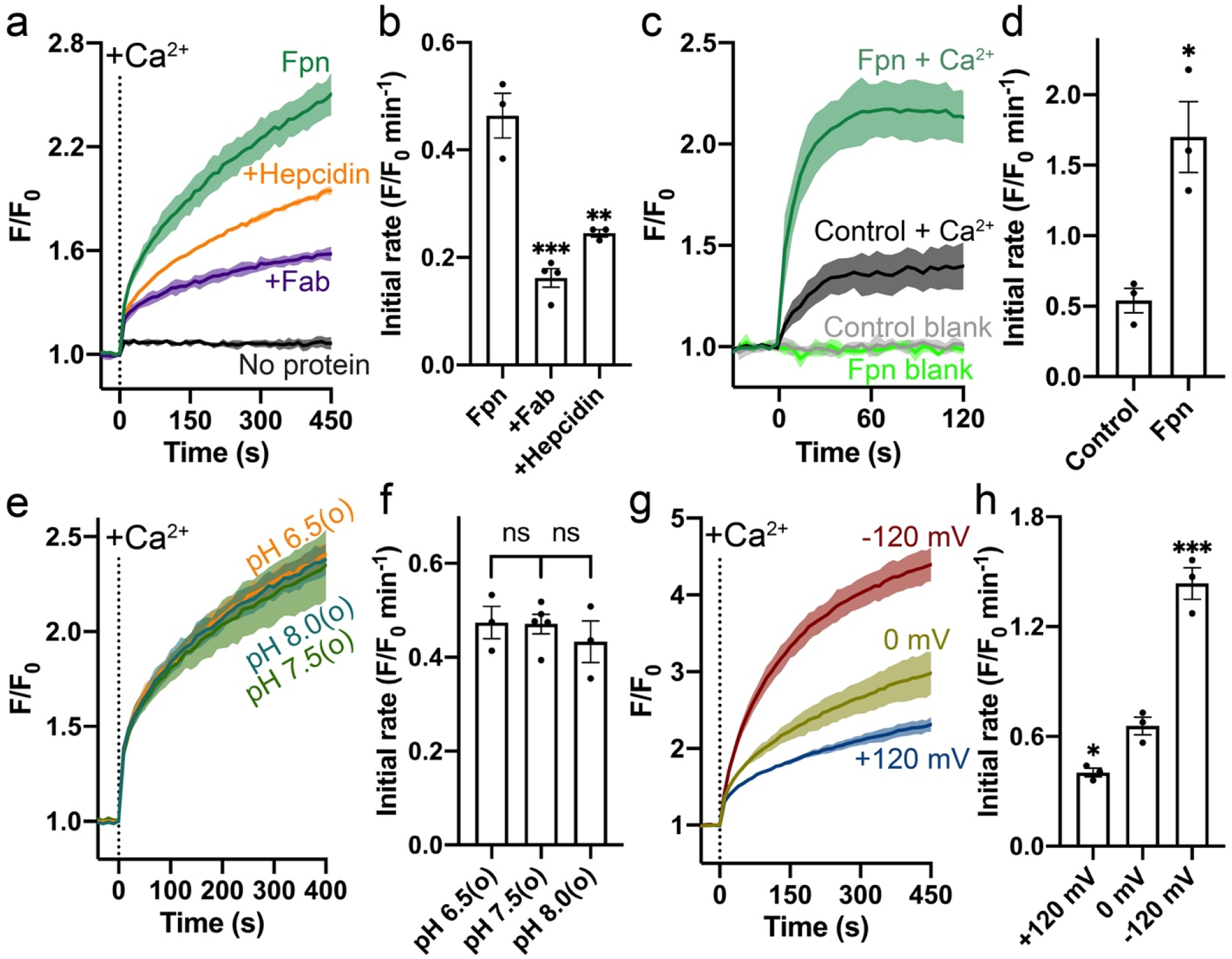
Specific Ca^2+^ uniport by HsFpn. (**a**) Ca^2+^ influx by HsFpn in proteoliposome measured by fluorescence changes (F/F_0_) of Fluo-4 (green). The presence of 11F9 Fab (purple) or hepcidin (orange) reduces the Ca^2+^ influx. (**b**) Comparison of initial rates of Ca^2+^ influx in (**a**). (**c**) Ca^2+^ uptake in HEK cells expressing HsFpn monitored by Fluo-4 loaded inside cells. Cells overexpressing Fpn show a significantly faster Ca^2+^ uptake compared to control cells transfected with an empty vector. (**d**) Comparison of initial rates of Ca^2+^ uptake in (**c**). (**e**) Ca^2+^ influx by HsFpn in proteoliposome at different outside (o) pHs. The inside pH is maintained at 7.5. (**f**) Comparison of initial rates of Ca^2+^ transport at different outside pHs. (**g**) Ca^2+^ influx by HsFpn in proteoliposome at different membrane potentials. Valinomycin was used to generate a membrane potential prior to the addition of Ca^2+^. (**h**) Comparison of initial rates of Ca^2+^ transport at different membrane potentials. In all the transport assays, 500 µM of Ca^2+^ was added at time zero. Statistical significances were analyzed with one-way analysis of variance (ANOVA) followed by Dunnett’s test for multiple comparisons. In this article, all time-dependent fluorescence traces are shown as solid lines (mean) with shaded regions (standard deviation, SD) from at least three repeats. For all bar graphs, a scatter plot is overlaid on each bar. The height represents the mean of at least three measurements, and the error bar standard error of the mean (SEM). Statistical significances are indicated: ns, not significant; *, *p*□<□0.05; **, *p*LJ<LJ0.01; ***, *p*□<□0.001; ****, *p* < 0.0001.

To further understand the mechanism of Ca^2+^ transport by Fpn, we examined whether common ions, H^+^, Na^+^, K^+^, and Cl^-^, are involved or required. Unlike the Fe^2+^ transport, the Ca^2+^ transport by Fpn is not coupled to H^+^ transport (**Figure 1e–f** and **Figure 1—figure supplement 2**). The absence of Na^+^, K^+^, or Cl^-^ also has no significant effect on Ca^2+^ transport (**Figure 1—figure supplement 3**). These results suggest that Fpn is a Ca^2+^ uniporter. If it is the case, Ca^2+^ influx into proteoliposomes would build up a positive membrane potential that hinders further import of Ca^2+^. We tested this in the following experiments. First, we measured Ca^2+^ transport at defined membrane potentials, +120 mV, 0 mV, and -120 mV, established by a K^+^ gradient in the presence of valinomycin, a K^+^ selective ionophore (**Figure 1g**). As shown in **Figure 1g–h**, Ca^2+^ transport is significantly enhanced at -120 mV and reduced at +120 mV, and Ca^2+^ transport at 0 mV is larger than that in the absence of a clamped membrane potential. These results are consistent with HsFpn being a Ca^2+^ uniporter.

Second, we measured Ca^2+^ transport at different concentrations of Ca^2+^ ([Ca^2+^]), first in the absence of a preset membrane potential and then in the presence of -120 mV membrane potential (**Figure 1—figure supplement 4a**). We first monitored Ca^2+^ influx in the presence of a 100-fold K^+^ gradient but the absence of valinomycin. After the influx reaches a steady state (∼460 sec), we added valinomycin to activate the membrane potential and we observed a further large increase of Ca^2+^ influx, and the rate of Ca^2+^ transport in the second phase is faster than that in the first phase. These results are consistent with HsFpn being a Ca^2+^ uniporter. The *K*_M_ for Ca^2+^ transport, calculated based on the initial rates of the second phase, is 48.5 (27.3 – 88.8) µM (95% confidence interval in parentheses) (**Figure 1—figure supplement 4b**). We measured Ca^2+^ uptake in HEK cells expressing HsFpn, and observed similar behavior of HsFpn in transporting Ca^2+^ (*K*_M_ = 85.9 (36.6 – 193.0) µM, **Figure 1—figure supplement 4c–d**). Although the transport rate initially increases linearly with increasing [Ca^2+^], it levels off with further increases of [Ca^2+^]. This behavior is consistent with Fpn being a Ca^2+^ transporter but not a Ca^2+^ channel.

### Cryo-EM structure of HsFpn bound to Ca^2+^

We determined the structure of HsFpn-11F9 in the presence of 2 mM Ca^2+^ by cryo-electron microscopy (cryo-EM) to an overall resolution of 3.0 Å (**Figure 2a** and **Figure 2—figure supplement 1**). The density map reaches a resolution of ∼2.4 Å in the TM regions and allows for the building and refinement of a structural model that contains residues 15 – 240, 289 – 400, and 452 – 555, which covers all 12 transmembrane (TM) helices (**Figure 2—figure supplement 2**). Residues 1 – 14, 241 – 288, 401 – 451, and 556 – 571, which are predicted to be disordered regions located to either the N- or C-terminus or loops between TM helices, are not resolved. The 12 TMs form two well-defined domains, with the N-terminal domain (NTD) composed of TM1-6 and the C-terminal domain (CTD) TM7-12 (**Figure 2b**). In the density map (**Figure 2a**), we noticed a non-protein density corralled by four helices (TM1, TM3, TM4, and TM6) in the NTD, and assigned it as a Ca^2+^ ion. This density was not present in structures of Co^2+^-bound Fpn (Pan et al. 2020; Billesbolle et al. 2020). The side chains of five conserved residues, Asp39, Gln99, Asn100, Asn212, and Glu219, are within 4 Å of the Ca^2+^ (**Figure 2b–c**). While in structures of Co^2+^-bound HsFpn (manuscript submitted together), the side chains of Asp39 and His43 at S1 point towards the central cavity between the NTD and CTD, the side chain of Asp39 in the current structure points towards the interior of the four-helical bundle to coordinate Ca^2+^ (**Figure 2d**). His43 also assumes a different rotamer conformation in the current structure (**Figure 2d**). When the NTD and CTD are aligned individually with their counterparts in the Co^2+^-bound HsFpn, the NTD has a root mean squared distance (RMSD) of 1.32 Å and the CTD 0.51 Å. The larger RMSD in the NTD is due to deviations of the extracellular halves of the TM helices and the extracellular loop between TM3-4 (**Figure 2e–g**). The current structure assumes an outward-facing conformation in which the NTD and CTD make contact on the intracellular side. The overall conformation is similar to that of the Co^2+^-bound HsFpn (**Figure 2e–g** and **Figure 2—figure supplement 3b**). The bound Ca^2+^ is not solvent-accessible in the current structure.

**Figure 2.**
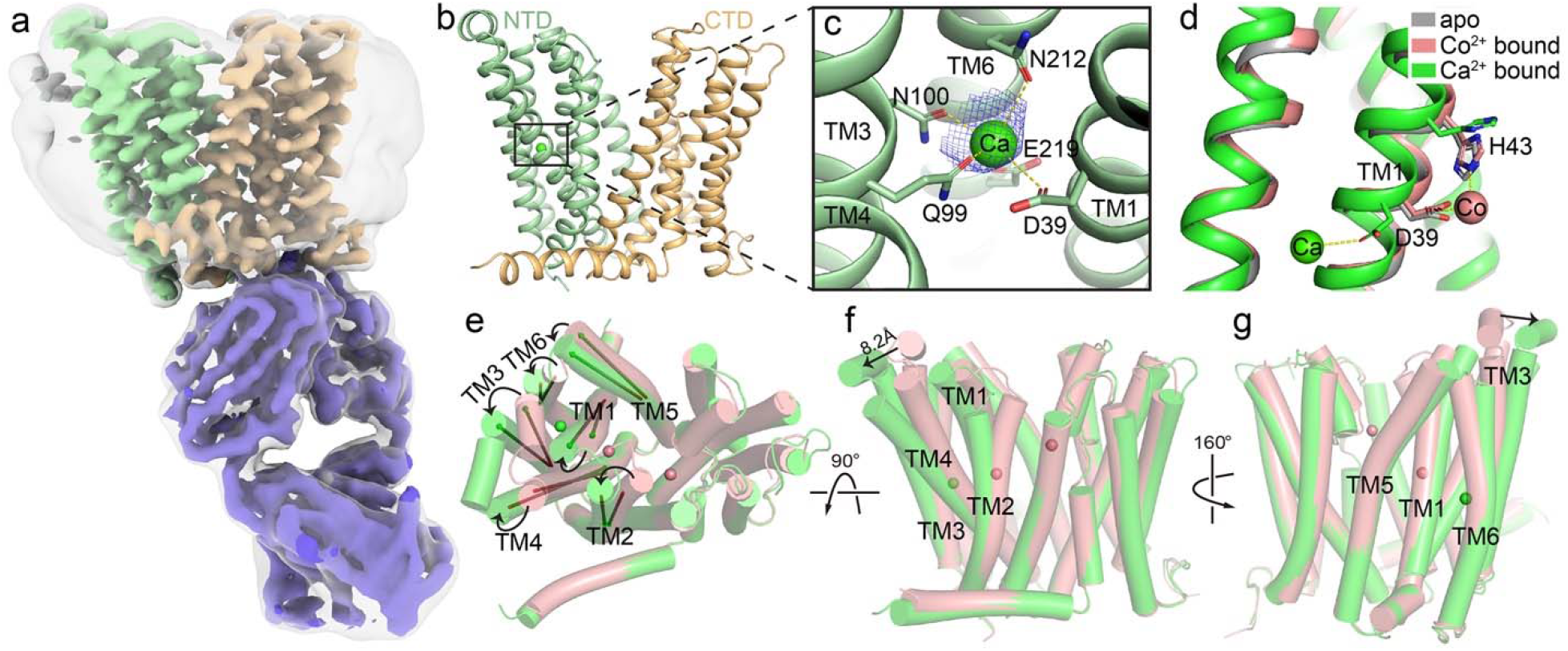
Structure of HsFpn bound to Ca^2+^. (**a**) Cryo-EM map of HsFpn in complex with 11F9 in the presence of Ca^2+^. Densities for NTD, CTD, and 11F9 are colored in pale green, light orange, and slate, respectively. A smoothened map contoured at a low threshold (translucent grey) is overlaid to show the lipid nanodisc density around Fpn. (**b**) Structure of HsFpn with a bound Ca^2+^ in an outward-facing conformation. NTD and CTD are colored as described in (**a**). Ca^2+^ is shown as a green sphere. (**c**) Zoomed-in view of the Ca^2+^ binding site in the NTD. The five residues coordinating Ca^2+^ are labeled and shown as side chain sticks. The density for Ca^2+^ is contoured at 7.5σ as blue mesh. (**d**) Comparison of apo (grey, PDB ID 6W4S), Co^2+^-bound (pink), and Ca^2+^-bound (green) HsFpn structures near the S1. The side chains of D39 and H43 are shown as sticks. Co^2+^ is shown as a pink sphere. (**e**), (**f**), and (**g**) Three views of conformational changes in NTD induced by Ca^2+^ binding. The Co^2+^-bound (pink) and Ca^2+^-bound (green) structures are aligned and shown as cartoon with cylindrical helices. (**e**) Top view (from the extracellular side) of the alignment. The helical directions of TM1-6 are visualized by vectors inside cylinders, and bending of these helices is indicated by black arrows. Bending of TMs viewed from the front (**f**) and the back (**g**). The displacement of the extracellular loop between TM3 and TM4 is marked with a black arrow and distance.

### Validation of the Ca^2+^ binding site

We validated the Ca^2+^ binding site by mutational studies. Both a single alanine mutation (Asp39Ala) and a triple alanine mutation (Gln99_Asn100_Glu219 to Ala) to residues at the Ca^2+^ binding site show significantly reduced Ca^2+^ transport activity (**Figure 3a–b**). In contrast, the double alanine mutation to S2 (Cys326Ala_His507Ala) does not affect Ca^2+^ transport significantly (**Figure 3a–b**). We also measured Ca^2+^ uptake by HsFpn mutants expressed in HEK cells (**Figure 3—figure supplement 1**) and found that alanine mutations to all five residues at the Ca^2+^ binding site significantly impair Ca^2+^ transport activity, while mutation to His43 (at S1), which is one helical turn away from the Ca^2+^ binding site, maintains high Ca^2+^ transport activity (**Figure 3c–d**).

**Figure 3.**
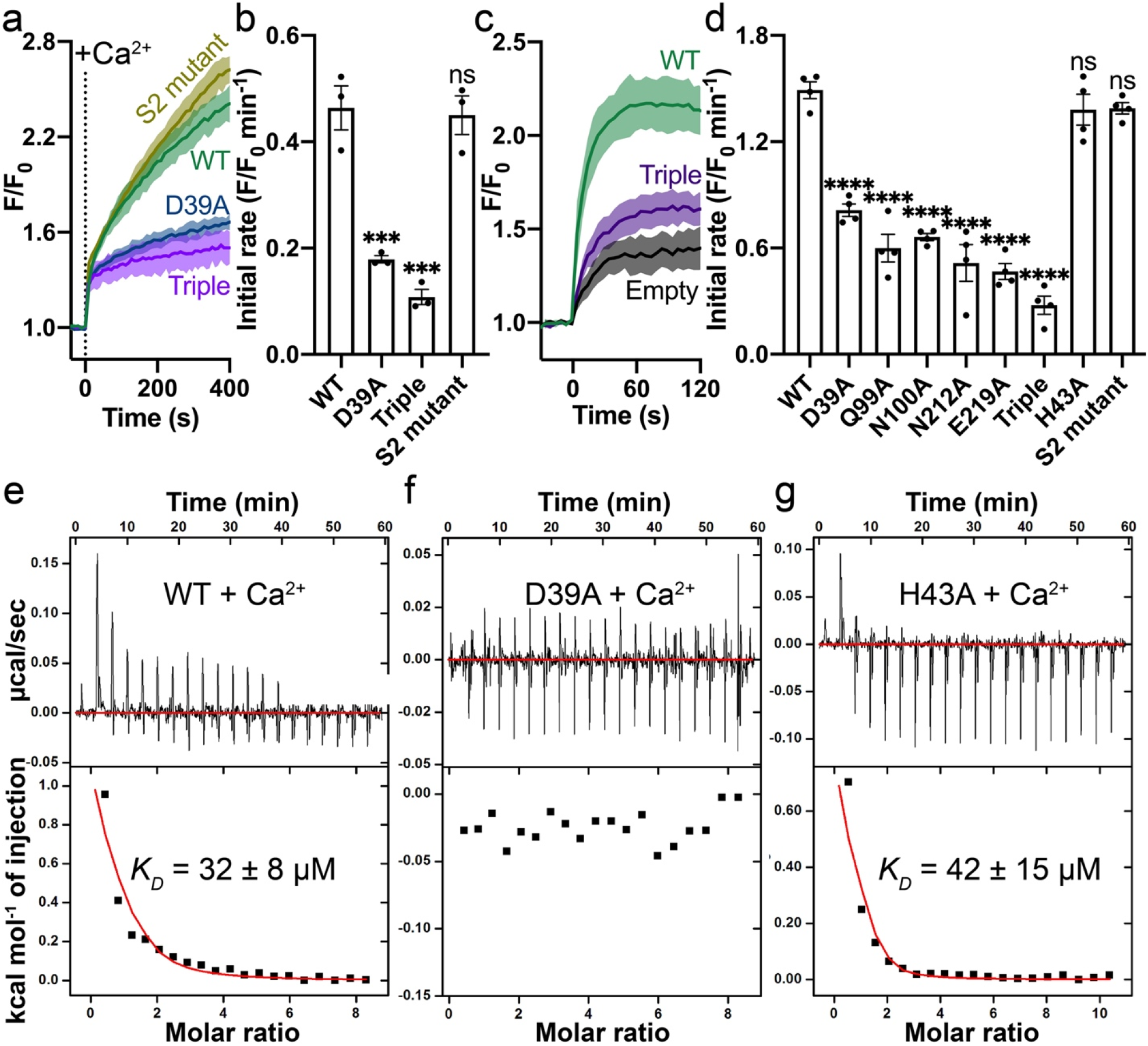
Mutations on the Ca^2+^ binding site. (**a**) Ca^2+^ influx by WT (green), D39A (blue), and the triple mutant (Q99A_N100A_E219A, purple) of the Ca^2+^ binding site in proteoliposome. (**b**) Comparison of initial rates of Ca^2+^ influx in (**a**). (**c**) Ca^2+^ uptake in HEK cells expressing WT (green) or the triple mutant (purple) Fpn. (**d**) Fpn-specific Ca^2+^ transport activities of WT and mutants. Initial rates are subtracted from the empty control. For statistical significances in (**b**) and (**d**), Dunnett’s test was used as a post hoc test following one-way ANOVA with the WT as the control. 500 µM Ca^2+^ was used in (**a**) – (**d**). (**e**), (**f**), and (**g**) Binding of Ca^2+^ to WT, D39A, and H43A HsFpn measured by ITC. Upper plot: raw thermogram showing the heat during binding and baseline (red line). Lower plot: integrated heat of each injection (black square) and the fit of data (red line).

We then measured the binding affinity of Ca^2+^ by isothermal titration calorimetry (ITC), and found that Ca^2+^ binding is an endothermic process with a dissociation constant (*K*_*D*_) of ∼32 µM (**Figure 3e**). Similar endothermic binding of Ca^2+^ was also reported in mouse Fpn and a bacterial homolog of Fpn (Deshpande et al. 2018). The 11F9 Fab does not interfere with Ca^2+^ binding as the HsFpn-11F9 complex has a similar Ca^2+^ binding affinity (*K*_*D*_ = 20 ± 6 µM) (**Figure 3— figure supplement 2a**).

We then measured Ca^2+^ binding to HsFpn mutants. We expressed and purified HsFpn with alanine mutations to the five residues at the Ca^2+^ binding site, and found that Ca^2+^ binding is abolished in these mutants (**Figure 3f** and **Figure 3—figure supplement 2e–h**). As a control, we also measured Ca^2+^ binding to His43Ala, which is one helical turn away from Asp39, and to the S2 mutant, and these mutants retain Ca^2+^ binding with affinities similar to that of the wild-type (WT) (**Figure 3g** and **Figure 3—figure supplement 2c**). Similarly, the alanine mutation to Gln99, which is part of the Ca^2+^ binding site, does not significantly change its Co^2+^ binding affinity (*K*_*D*_ = 226 ± 23 µM) (**Figure 3—figure supplement 2d**). These results provide further support to the observed Ca^2+^ binding site in HsFpn.

### Competition between Ca^2+^ and Co^2+^ in HsFpn

Next, we measured the binding and transport of Ca^2+^ in the presence of Co^2+^ or Fe^2+^, and *vice versa*. In the presence of 2 mM Co^2+^, Ca^2+^ binding affinity is reduced by ∼4-fold as shown in **Figure 4a**. The reduced binding is apparent both from the reduced heat absorption during titration, and the reduced *K*_*D*_ from the overall fitting of the ITC data. Reduced Ca^2+^ binding in the presence of Co^2+^ is consistent with the structures due to the shared Asp39 by the Ca^2+^ binding site and S1 for Co^2+^ (**Figure 2d**). On the other hand, Co^2+^ binding affinity (*K*_*D*_) is not affected in the presence of 2 mM Ca^2+^, as shown in **Figure 4b**, although the amount of heat release is appreciably less in the presence of Ca^2+^. This result is in line with the effect of the S1 mutation(Asp39Ala) on Co^2+^ binding (Pan et al. 2020), as competition between Ca^2+^ and Co^2+^ at S1 does not affect transition metal ion binding at S2.

We then measured Ca^2+^ transport in the presence of Fe^2+^ or Co^2+^, and Fe^2+^ or Co^2+^ transport in the presence of Ca^2+^. We tested conditions in which Ca^2+^ and Fe^2+^ are either on the opposite or the same side of the membrane (**Figure 4c** and **Figure 4—figure supplement 2a–b**). We found that while Fe^2+^ or Co^2+^ transport is not sensitive to the presence of Ca^2+^ (**Figure 4c–d** and **Figure 4—figure supplement 1a–b**), Ca^2+^ transport is significantly reduced in the presence of Fe^2+^ or Co^2+^ (**Figure 4e–g** and **Figure 4—figure supplement 2c–e**). This is consistent with our understanding that the Ca^2+^ transport in Fpn is mediated by a single site that is significantly affected by the presence of Fe^2+^, while the Fe^2+^/2H^+^ exchange is mediated by both S1 and S2 and that the Ca^2+^ binding only disturbs S1. In the presence of Fe^2+^, competition at S1 would hamper Ca^2+^ binding and transport, while in the presence of Ca^2+^, the entire S2 and His43 of S1 remain available and hence allow the Fe^2+^/2H^+^ exchange (**Figure 4—figure supplement 3a–c**). Further studies are needed to dissect the intertwined transport pathways of Fe^2+^ and Ca^2+^ transport in Fpn. We conclude that HsFpn functions mainly as a Fe^2+^/2H^+^ exchanger in the presence of Fe^2+^, however, in the absence of Fe^2+^, Fpn mediates Ca^2+^ transport.

**Figure 4.**
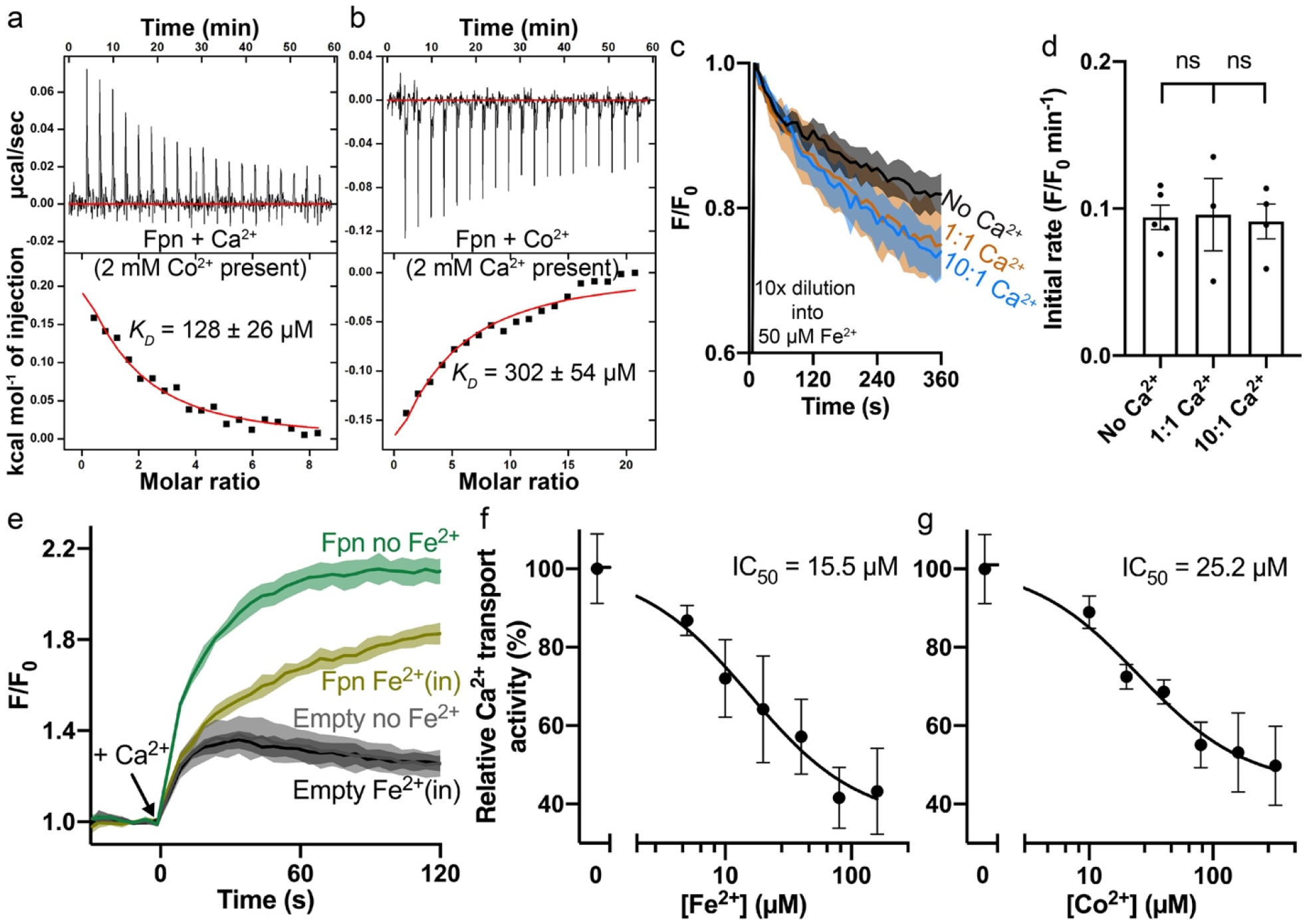
Interplay between Fe^2+^/Co^2+^ and Ca^2+^ in HsFpn. (**a**) Ca^2+^ binding in the presence of 2 mM Co^2+^. (**b**) Co^2+^ binding in the presence of 2 mM Ca^2+^. (**c**) Fe^2+^ influx by HsFpn in the presence of Ca^2+^ in proteoliposome. Fe^2+^ influx was initiated by 10-fold (10×) dilution of liposome samples into outside buffers containing 50 µM Fe^2+^. The “1:1” or “10:1” denotes the ratio of [Ca^2+^] inside:outside liposomes achieved after 10× dilution. 1 mM sodium ascorbate was included in the samples. All fluorescence traces are subtracted from their corresponding no protein controls at the same conditions. (**d**) Comparison of initial rates of Fe^2+^ transport in (**g**). One-way ANOVA was used for statistical analysis. (**e**) Ca^2+^ (1 mM) uptake in HEK cells expressing HsFpn in the presence of Fe^2+^ (80 µM) loaded inside cells (dark yellow) or in the absence of Fe^2+^ (green) monitored by jGCaMP7s co-expressed in cytosol. Cells transfected with an empty vector (dark and light gray) serve as negative controls. (**f**) and (**g**) Reduced Ca^2+^ transport at different concentrations of Fe^2+^ and Co^2+^. The relative Ca^2+^ transport activities represent the normalized initial rates of Fpn-specific Ca^2+^ uptake. The IC_50_ values were calculated from curve fits (black solid line) to an inhibitory dose-response equation.

## Discussion

In summary, we visualize the Ca^2+^ binding site in HsFpn, and we show that Ca^2+^ transport by Fpn follows a uniport mechanism and is different from that of Fe^2+^ transport. While Fe^2+^ export is coupled to the import of H^+^ and is mediated by both S1 and S2, Ca^2+^ import is mediated by a single binding site and is not coupled to another ion (**Figure 1e–f** and **Figure 1—figure supplement 2**–**3**). We also show that the *K*_M_ for Ca^2+^ transport is ∼25-fold larger than that for Fe^2+^ transport, indicating that Fpn has a higher sensitivity to transition metal ions. The Ca^2+^ binding site in HsFpn is equivalent to a Ca^2+^ binding site previously identified in a bacterial homolog of Fpn (*Bdellovibrio bacteriovorus*; BbFpn) (**Figure 2—figure supplement 3a**) (Deshpande et al. 2018). Although the overall sequence identity between HsFpn and BbFnp is only 26%, residues at the Ca^2+^ binding site are conserved except for Asn100 in HsFpn, which is Phe85 in BbFpn.

Our discovery of Ca^2+^ transport in Fpn demonstrates a novel Ca^2+^ entry pathway in cells expressing Fpn. We emphasize that the Ca^2+^ transport by Fpn is sensitive to [Fe^2+^], and significant Ca^2+^ influx occurs when [Fe^2+^] is low (**Figure 4—figure supplement 3**). Our study also establishes Fpn as a transporter capable of operating with two different transport mechanisms. Since Ca^2+^ is an important second messenger to various physiological responses in cells, Ca^2+^ entry through Fpn provides a link between iron homeostasis and cellular responses.

## Materials and methods

### Cloning, expression, and purification of human Fpn (HsFpn)

The codon-optimized cDNA of HsFpn (UniProt ID: <u>Q9NP59</u>) was cloned into a pFastBac dual vector. A Tobacco Etch Virus (TEV) protease site and an octa-histidine (8×His) tag was appended to the C-terminus of the protein. HsFpn was expressed in Sf9 (*Spodoptera frugiperda*) cells using the Bac-to-Bac method (Invitrogen). Purification of HsFpn follows the same protocol reported for *Tarsius syrichta* Fpn (TsFpn) (Pan et al. 2020). Purified HsFpn was collected from size-exclusion chromatography (SEC) in FPLC buffer containing 20 mM HEPES, pH7.5, 150 mM NaCl, and 1 mM (w/v) n-dodecyl-β-D-maltoside (DDM, Anatrace). Mutations to HsFpn were generated using the QuikChange method (Stratagene) and verified by sequencing. Mutants were expressed and purified following the same protocol for the WT.

### Preparation of HsFpn-11F9 complex in nanodisc

Membrane scaffold protein (MSP) 1D1 was expressed and purified following an established protocol (Martens et al. 2016). For lipid preparation, 1-palmitoyl-2-oleoyl-sn-glycero-3-phospho-(1’-rac)-choline (POPC, Avanti Polar Lipids), 1-palmitoyl-2-oleoyl-sn-glycero-3-phospho-(1’-rac)-ethanolamine (POPE, Avanti Polar Lipids) and 1-palmitoyl-2-oleoyl-sn-glycero-3-phospho-(1’-rac)-glycerol (POPG, Avanti Polar Lipids) were mixed at a molar ratio of 3:1:1, dried under Argon and resuspended with 14 mM DDM (Autzen et al. 2018). For nanodisc reconstitution, HsFpn, MSP1D1 and lipid mixture were mixed at a molar ratio of 1:2.5:50 and incubated on ice for 1h. Detergents were removed by the sequential addition of 60 mg/mL Biobeads SM2 (Bio-Rad) for three times within 3 h. The sample was then incubated with Biobeads overnight at 4 °C. After removal of Biobeads, 11F9 was added to the sample at a molar ratio of 1.1:1 to HsFpn. The complex was incubated on ice for 30 min before being loaded onto a SEC column equilibrated with 20 mM HEPES, pH7.5, 150 mM NaCl. The purified nanodisc sample was concentrated to 10 mg/ml and incubated with 2 mM CaCl_2_ for 30 min on ice before cryo-EM grid preparation.

### Cryo-EM sample preparation and data collection

The cryo-EM grids were prepared using Thermo Fisher Vitrobot Mark IV. The Quantifoil R1.2/1.3 Cu grids were glow-discharged with air for 15 s at 10 mA using Plasma Cleaner (PELCO EasiGlow™). Aliquots of 3.5 µl nanodisc sample were applied to the glow-discharged grids. After blotted with filter paper (Ted Pella, Inc.) for 4.0 s, the grids were plunged into liquid ethane cooled with liquid nitrogen. A total of 4,498 micrograph stacks were collected on a Titan Krios at 300 kV equipped with a K3 direct electron detector (Gatan) and a Quantum energy filter (Gatan) at a nominal magnification of 81,000× and defocus values from -2.5 µm to -0.8 µm. Each stack was exposed in the super-resolution mode with an exposing time of 0.0875 s per frame for a total of 40 frames per stack. The total dose was about 50 e^-^/Å^2^ for each stack. The stacks were motion corrected with MotionCor2 (Zheng et al. 2017) and binned 2-fold, resulting in a pixel size of 1.10 Å/pixel. In the meantime, dose weighting was performed (Grant and Grigorieff 2015). The defocus values were estimated with Gctf (Zhang 2016).

### Cryo-EM data processing

A total of 2,184,301 particles were automatically picked based on a reference map of TsFpn-11F9 (EMD-21460) low-pass filtered to 20 Å in RELION 3.1 (Scheres 2015, 2012; Kimanius et al. 2016; Zivanov et al. 2018). Particles were extracted and imported to CryoSparc (Punjani et al. 2017) for 2D classification. A total of 1,329,782 particles were selected from good classes in 2D classification, which display recognizable structural features. Four 3D references were generated by *ab initio* reconstruction with limited particles from the best 2D classes. Multiple rounds of heterogeneous refinement were performed with particles selected from the 2D classification and four initial reference models until no more than 5% of input particles were classified into bad classes. A final of 437,959 particles after heterogeneous refinement were subjected to non-uniform (NU) refinement with an adaptive solvent mask. After handedness correction, local refinement and CTF refinement were performed with a soft mask around the Fpn and the Fv region of the Fab. Resolutions were estimated with the gold-standard Fourier shell correlation 0.143 criterion (Rosenthal and Henderson 2003). Local resolution of the maps was estimated in CryoSparc (Punjani et al. 2017).

### Model building and refinement

The structure of apo HsFpn (from PDB ID 6W4S) and Fab 11F9 (from PDB ID 6VYH) were individually docked into density maps in Chimera (Pettersen et al. 2004). The docked model was manually adjusted with added ligands in COOT (Emsley et al. 2010). Structure refinements were carried out by PHENIX in real space with secondary structure and geometry restraints (Adams et al. 2010). The EMRinger Score was calculated as described (Barad et al. 2015). All structure figures were prepared in Pymol and ChimeraX (Pettersen et al. 2021).

### Isothermal titration calorimetry

The WT and mutant HsFpn proteins were purified as described above and concentrated to 50 – 75 μM (3 – 4.5 mg/mL) in the FPLC buffer. The buffer was degassed, and all the protein samples were centrifuged at 18,000 × g for 20 min to remove aggregates. The injectant of 2 mM CaCl_2_ or 5 mM CoCl_2_ was prepared in the same FPLC buffer. For competition binding, either 2 mM CoCl_2_ or CaCl_2_ was added to protein samples prior to ligand titration. The ITC measurements were performed in Auto-iTC200 (MicroCal) at 25 °C. A total of 25 injections were administered (1.01 μL for injections 1 and 2.02 μL for injections 2 – 25) with a 150 s interval between injections. Background-subtracted data were fitted with binding models in the Origin 8 software package (MicroCal) to extract *K*_*D*_, ΔH, and entropy change (ΔS).

### Expression of HsFpn in HEK cells

The cDNAs of WT and mutant HsFpn were subcloned into a modified pEG BacMam vector with a C-terminal Strep-tag. The resulting plasmids with Fpn or the empty plasmid were transfected into HEK 293S cells on 6-well plates with 293fectin™ transfection reagent (Invitrogen/Thermo Fisher) per the manufacturer’s protocol. After incubation at 37 °C with 8% CO_2_ for 2 days, cells were harvested and solubilized in the lysis buffer (20 mM HEPES, pH 7.5, 150 mM NaCl, 10% glycerol) plus 1% LMNG and Protease Inhibitor Cocktail (Roche) for 1 h at 4°C. Insoluble fractions were pelleted by centrifugation and supernatants were run in SDS-PAGE. Proteins were visualized by western blotting with mouse anti-Strep (Invitrogen/Thermo Fisher) and IRDye-800CW anti-mouse IgG (Licor). Images were taken on an Odyssey infrared scanner (Licor).

### Ca^2+^ uptake and H^+^ transport assays in HEK cells

The pEG BacMam plasmids with HsFpn or the empty plasmid were transfected into HEK 293S cells on black wall 96-well microplates coated with poly-D-lysine (Invitrogen/Thermo Fisher). After 2 days, cells were washed in the live cell imaging solution (LCIS) containing 20 mM HEPES, 140 mM NaCl, 2.5 mM KCl, 1.0 mM MgCl_2_, 5 mM D-glucose, pH 7.4. The loading of Fluo-4 (Invitrogen/Thermo Fisher, AM, cell-permeant) for Ca^2+^ uptake assays or pHrodo™ Red (Invitrogen/Thermo Fisher, AM) for H^+^ transport assays was performed following the manufacturer’s protocols. After the loading finished, free dyes were washed away, and cells in each well were maintained in 90 µL LCIS. Both the Ca^2+^ uptake and H^+^ transport assays were performed in the FlexStation 3 Multi-Mode Microplate Reader (Molecular Devices) at 37 °C. Fluorescence changes were recorded at an excitation and emission wavelength of 485 nm and 538 nm for Ca^2+^ uptake assays, and 544 nm and 590 nm for H^+^ transport assays with 5 s intervals. Transport was triggered by the addition of 10 µL ligand stock solution (CaCl_2_ or CoCl_2_) to achieve the desired concentration of extracellular Ca^2+^ or Co^2+^. The H^+^ transport was assayed with 500 µM Co^2+^ or Ca^2+^. To test the effect of extracellular pH on Ca^2+^ uptake, the extracellular buffer was changed to pre-warmed LCIS with adjusted pH soon before the addition of 500 µM Ca^2+^. All the mutants were assayed with 500 µM Ca^2+^. For Ca^2+^ uptake assays, the slopes of straight lines fitted to transport data within 25 s were used to represent initial rates. For H^+^ transport assays, relative fluorescence changes at the equilibrium stage were averaged to represent intracellular pH changes.

### Ca^2+^ uptake in the presence of Fe^2+^ or Co^2+^ in HEK cells

The WT and mutant HsFpn were expressed in HEK 293S cells as described above except that the plasmid for mammalian expression of jGCaMP7s (pGP-CMV-jGCaMP7s) was co-transfected. The pGP-CMV-jGCaMP7s was a gift from Douglas Kim & GENIE Project (Addgene plasmid # 104463; http://n2t.net/addgene:104463; RRID:Addgene_104463) (Dana et al. 2019). Ca^2+^ uptake assays in the presence of Fe^2+^ or Co^2+^ were performed in FlexStation 3 similarly as described above except that the excitation and emission wavelength were set at 485 nm and 513 nm. Fe^2+^ (NH_4_Fe(SO_4_)_2_) or Co^2+^(CoCl_2_) ions were first loaded into cells for >5 min at 37 °C, while fluorescence readings were recorded. For Fe^2+^ loading, 1 mM sodium ascorbate was used to protect Fe^2+^ ions from oxidation. To start the export of Fe^2+^ or Co^2+^, the extracellular buffer was exchanged to pre-warmed LCIS soon before the addition of Ca^2+^. LCIS with adjusted pH was used when testing the effect of extracellular pH. Straight lines were used to fit transport data within 25 s, and their slopes were used to represent initial rates.

### Reconstitution of HsFpn into liposomes

POPE and POPG lipid (Avanti Polar Lipids) were mixed at a 3:1 molar ratio, dried under Argon, and vacuumed overnight to remove chloroform. The dried lipid was resuspended in reconstitution buffer (20 mM HEPES, pH 7.5, 100 mM KCl) to a final concentration of 10 mg/mL. After hydration for 2 h, the liposome sample was sonicated to transparency and incubated with 40 mM n-decyl-β-D-maltoside (DM, Anatrace) for 2 h at room temperature under gentle agitation. Then HsFpn protein was added at a 1:100 (w/w, protein:lipid) ratio. For the empty control, the same volume of blank buffer was added. Detergent was removed by dialysis at 4 °C against the reconstitution buffer. Dialysis buffer was changed every day for 4 days. The proteoliposome or empty liposome sample was aliquoted and frozen with liquid nitrogen, and was stored at -80 °C for future use.

### Ca^2+^ and Fe^2+^/Co^2+^ influx assays in proteoliposomes

Proteoliposomes with HsFpn or empty liposomes were thawed and mixed with 100 μM Fluo-4 (Invitrogen/Thermo Fisher, cell impermeant) for Ca^2+^ influx assays, or 250 μM calcein (Invitrogen/Thermo Fisher) for Co^2+^/Fe^2+^ influx assays. The dye was incorporated during three cycles of freeze-thaw. Liposomes were extruded to homogeneity with a 400 nm filter (NanoSizer™ Extruder, T&T Scientific Corporation). Removal of free dyes outside liposomes and exchange of outside buffer was achieved by passing samples through a desalting column (PD-10, GE Healthcare) equilibrated with the outside buffer. Liposome samples were transferred to a quartz cuvette for fluorescence recording in a FluoroMax-4 spectrofluorometer (HORIBA). Fluorescence changes were recorded at an excitation and emission wavelength of 494 nm and 513 nm with 10 s intervals at 37 °C.

For Ca^2+^ influx assays, transport was initiated by the addition of CaCl_2_ to the desired concentration. When testing the inhibition by hepcidin or 11F9 Fab, 20 µM human hepcidin (Sigma) or purified 11F9 Fab (Pan et al. 2020) was added prior to freeze-thaw cycles, and the transport was assayed with 500 µM CaCl_2_. When testing the effect of different pHs, outside buffers of the same components as the inside buffer but with adjusted pHs were used. When testing the effect of different common cations and anions, the inside and outside of liposomes have symmetrical buffers with 100 mM NaCl, KCl, or K-Gluconate. When testing the concentration-dependent electrogenic transport, the outside buffer containing 20 mM HEPES, pH 7.5, 1 mM KCl, and 99 mM NaCl was used, and 40 nM valinomycin was added after ∼460 s to clamp intravesicular potential at ∼-120 mV. When testing the effects of different membrane potentials, valinomycin was incubated with liposome samples for 5 min prior to the addition of CaCl_2_. For the -120 mV group, the outside buffer contained 20 mM HEPES, pH 7.5, 1 mM KCl and 99 mM NaCl. For the 0 mV group, the outside buffer was the same as the inside buffer. For the +120 mV group, the reconstitution buffer with NaCl (20 mM HEPES, pH 7.5, 1 mM KCl, 99 mM NaCl) was used during liposome preparation, and the outside buffer contained 100 mM KCl.

For Fe^2+^ or Co^2+^ influx assays, transport was initiated by the dilution of 30 µL of liposome sample into 270 µL of outside buffer containing 50 µM of NH_4_Fe(SO_4_)_2_ or CoCl_2_. In the case of Fe^2+^ influx, 1 mM sodium ascorbate was added. To load liposomes with Ca^2+^, 500 µM CaCl_2_ was added prior to freeze-thaw cycles. Buffer with 500 µM CaCl_2_ was used during the desalting step. To create a 10-fold Ca^2+^ gradient opposite to the Fe^2+^ or Co^2+^gradient, samples were diluted into the outside buffer without Ca^2+^.

## Data availability

The cryo-EM density map of nanodisc-encircled HsFpn-11F9 in the presence Ca^2+^ has been deposited in the Electron Microscopy Data Bank (https://www.ebi.ac.uk/pdbe/emdb/) under accession code EMD-27497. The corresponding atomic coordinate file has been deposited in the Protein Data Bank (http://www.rcsb.org) under ID code 8DL6. Any additional information required to reanalyze the data reported in this paper is available from the lead contact Yaping Pan upon request.

## Acknowledgments

This work was supported by grants from NIH (HL157473 to Y.P. and DK122784, HL086392 to M.Z.), and Cancer Prevention and Research Institute of Texas (R1223 to M.Z.). We acknowledge the use of the cryo-EM core at Baylor College of Medicine (BCM) for grid preparation and screening. Cryo-EM data in this work were acquired at the Stanford-SLAC Cryo-EM Center (S^2^C^2^) supported by the NIH Common Fund Transformative High Resolution Cryo-Electron Microscopy program (U24 GM129541), and at the Pacific Northwest Center for Cryo-EM (PNCC) at Oregon Health & Science University, supported by the NIH grant U24GM129547. We acknowledge L. Wang for help with grid preparation, and Z. Ren for making some of mutants. We are grateful to A. A. R. Adeosun for her instructions on the use of FlexStation 3.

## Author Contributions

M.Z., Y.P., and J.S. conceived the project. J.S., Y.P., and A.S.W. conducted experiments. Y.P. and J.S. wrote the initial draft of the manuscript, and all authors contributed to revisions of the manuscript.

---

**Figure 1—figure supplement 1.**
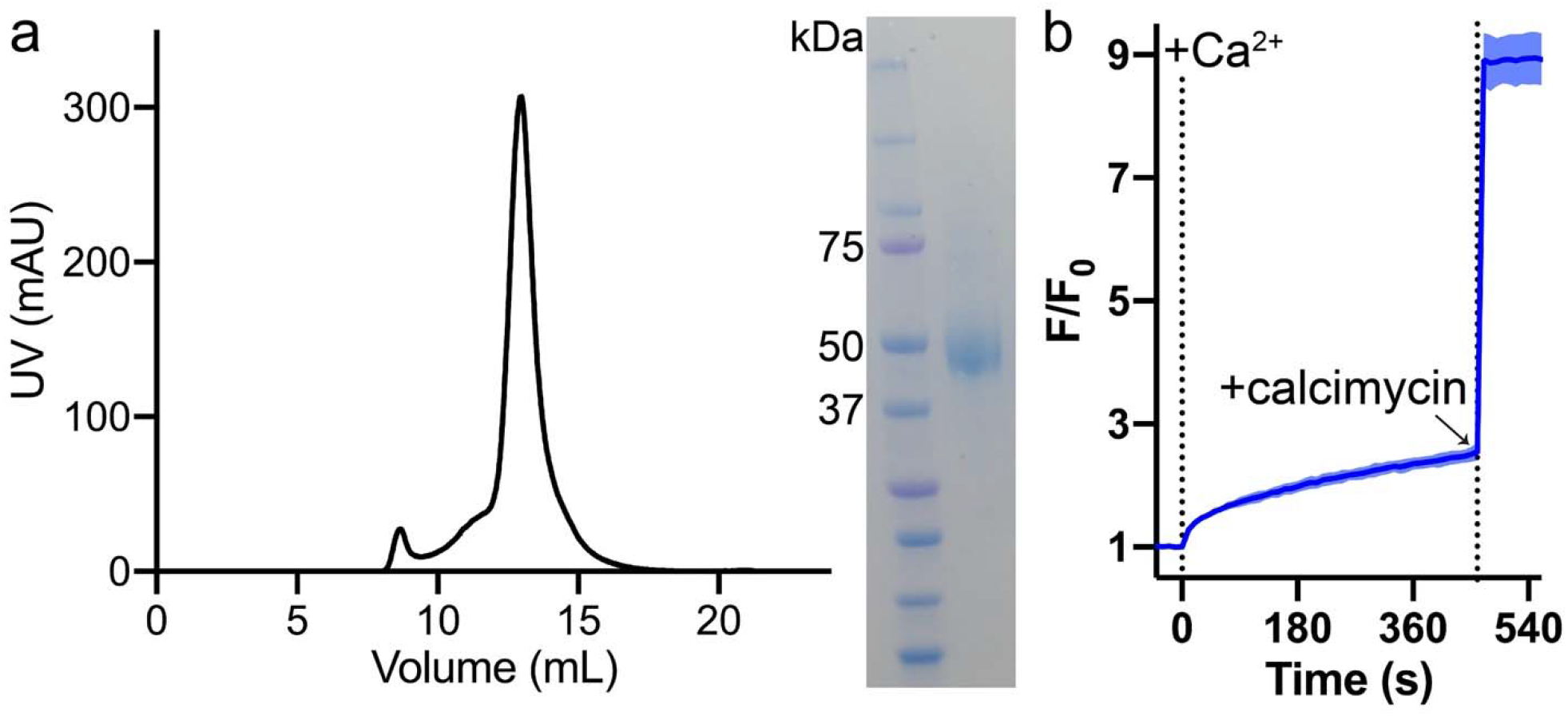
Expression of HsFpn and validation of Ca^2+^ transport in proteoliposome. (**a**) SEC profile (left) and SDS-PAGE gel image of purified HsFpn. (**b**) Calcimycin, a Ca^2+^ ionophore, could collapse the Ca^2+^ gradient after 450 s of Ca^2+^ influx by Fpn in proteoliposomes. 500 µM of Ca^2+^ was added at time zero.

**Figure 1—figure supplement 2.**
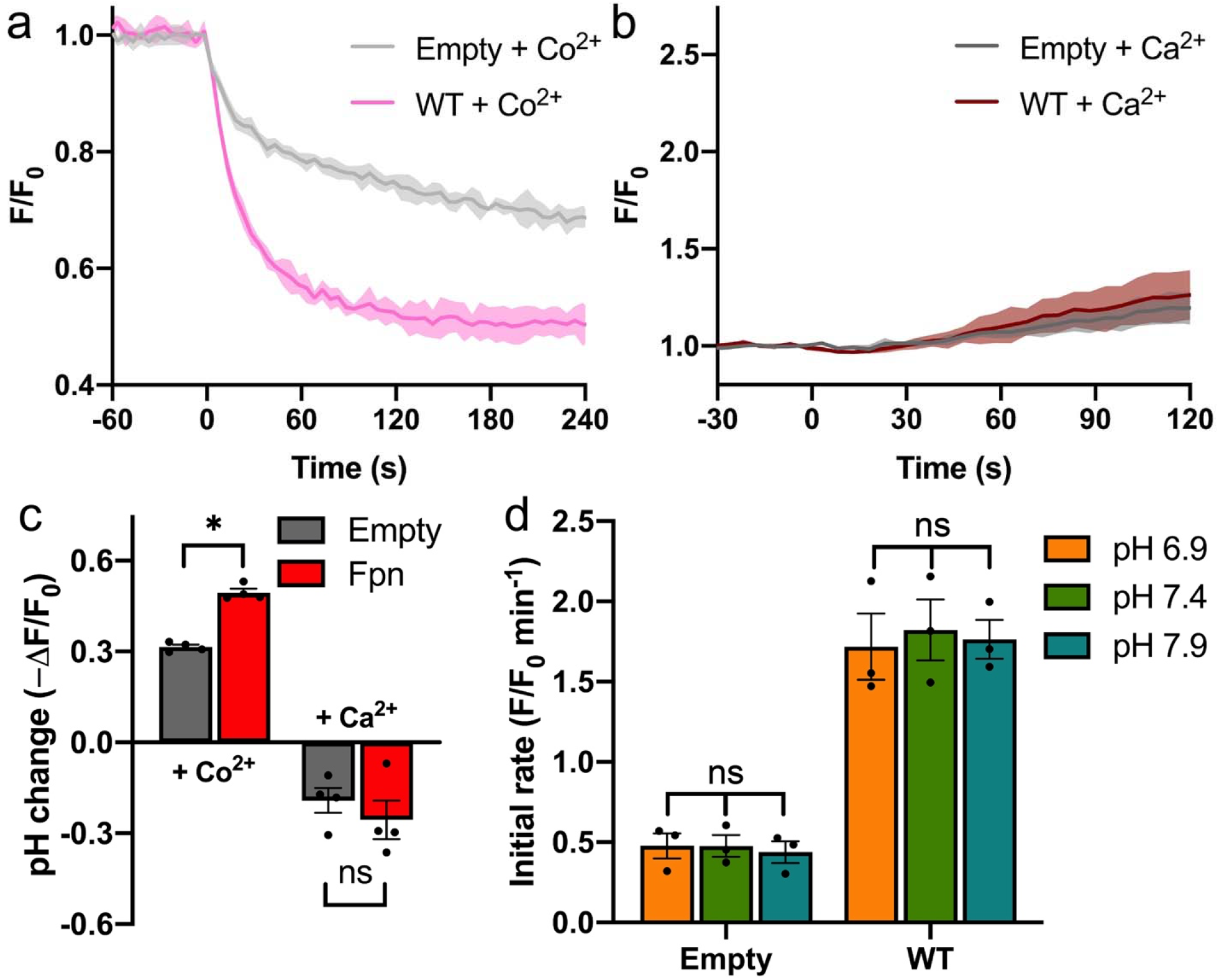
H^+^-independent Ca^2+^ uptake by HsFpn in HEK cells. (**a**) and (**b**) H^+^ transport induced by the Co^2+^ or Ca^2+^ uptake of Fpn monitored by a pH-sensitive dye (pHrodo Red). (**c**) Comparison of cytosolic pH changes induced by the Co^2+^ or Ca^2+^ uptake. The averaged final fluorescence changes from the traces in (**a**) and (**b**) are plotted. Decreased relative fluorescence (ΔF/F_0_) correlates to increased pH. Sidak’s multiple comparisons test was used following two-way ANOVA. (**d**) Ca^2+^ transport rates at different extracellular pHs. Two-way ANOVA: among different pHs, *p* = 0.912; between empty and WT, *p* < 0.0001; interaction, *p* = 0.918.

**Figure 1—figure supplement 3.**
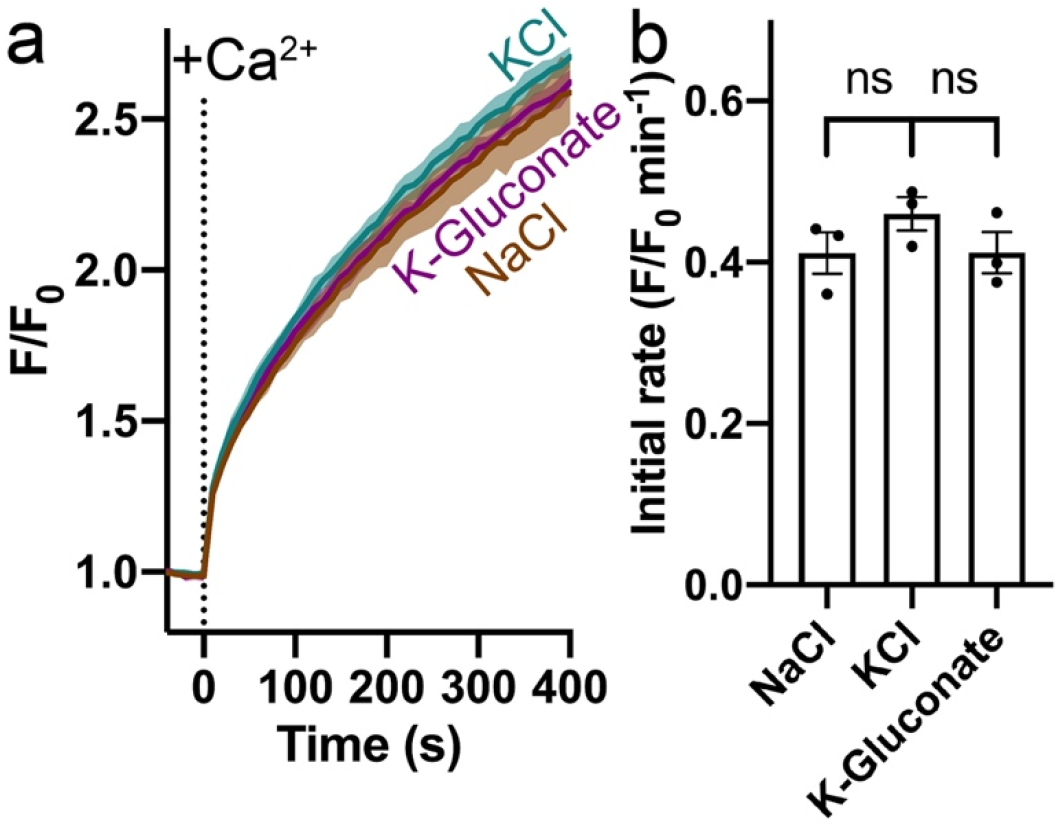
Effect of Na^+^, K^+^, and Cl^-^ on Ca^2+^ transport by HsFpn. (**a**) Ca^2+^ influx in proteoliposome with symmetrical NaCl, KCl, and K-Gluconate. 500 µM Ca^2+^ was added at time zero. (**b**) Comparison of initial rates of Ca^2+^ transport in different salts. Statistical significances were analyzed with one-way ANOVA.

**Figure 1—figure supplement 4.**
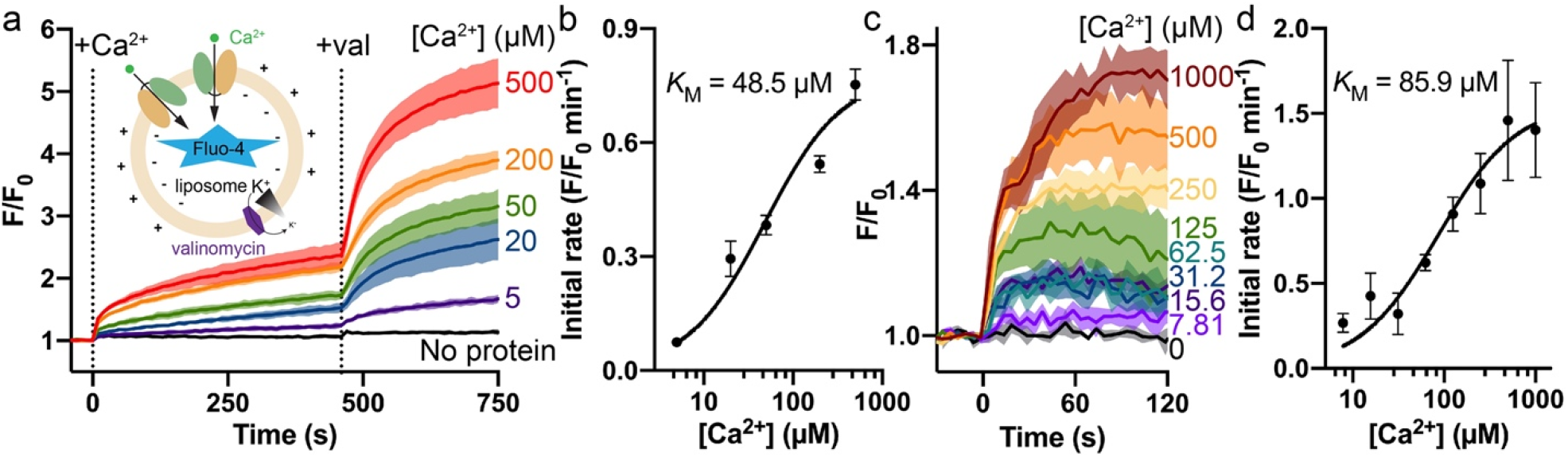
Ca^2+^ transport by Fpn. (a) Ca^2+^ transport into liposome with purified Fpn monitored by impermeable Fluo-4. Different concentrations of Ca^2+^ (labeled to the right of each trace) were added followed by the addition of valinomycin (val) to clamp membrane potential. The time points for the addition of Ca^2+^ or val are indicated by vertical dash lines. For the “No protein” control, 500 µM Ca^2+^ was used. The inset shows the schematic of the proteoliposome with a 100:1 inside:outside K^+^ gradient. (**b**) Initial rates of fluorescence change after the addition of val versus concentrations of Ca^2+^ for data in (**a**). (**c**) Dose-dependent Fpn-specific Ca^2+^ uptake in HEK cells expressing Fpn measured by Fluo-4 loaded inside cells. The raw traces of the empty control group at a certain [Ca^2+^] were subtracted from the corresponding traces of the Fpn WT group. (**d**) Initial rates of fluorescence change versus concentrations of Ca^2+^ in HEK cells. The black solid lines in (**b**) and (**d**) are curve fits to a Michaelis–Menten equation.

**Figure 2—figure supplement 1.**
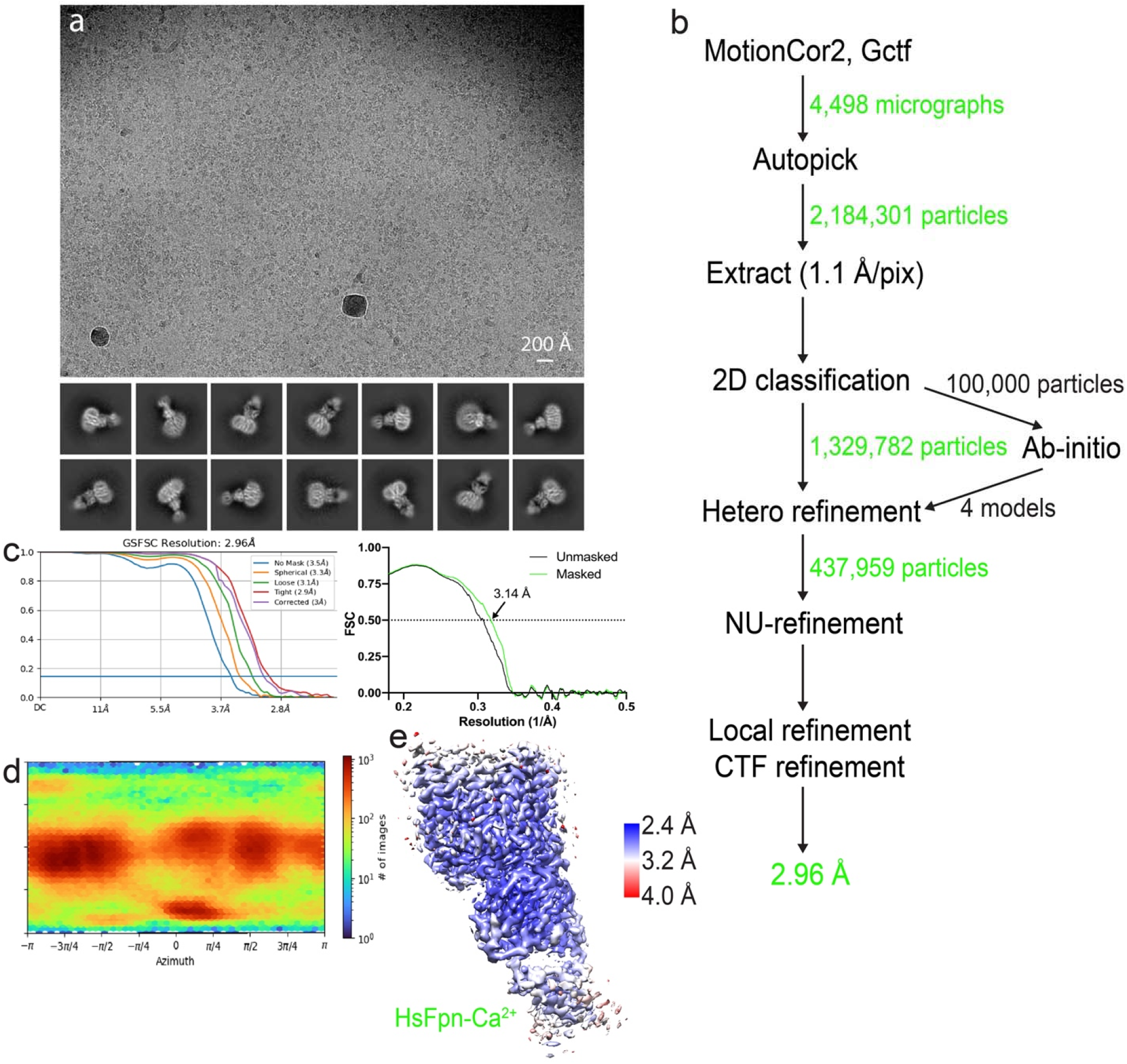
Cryo-EM analysis of HsFpn-Ca^2+^ in nanodisc. (**a**) Representative electron micrograph (upper panel) and 2D class averages (lower panel). (**b**) Workflow of single-particle data processing. (**c**) The gold-standard Fourier shell correlation (FSC) curves (left panel) and map-to-model FSC curves (right panel). (**d**) Direction distribution of particles used in the final 3D reconstruction. (**e**) Local resolution map.

**Figure 2—figure supplement 2.**
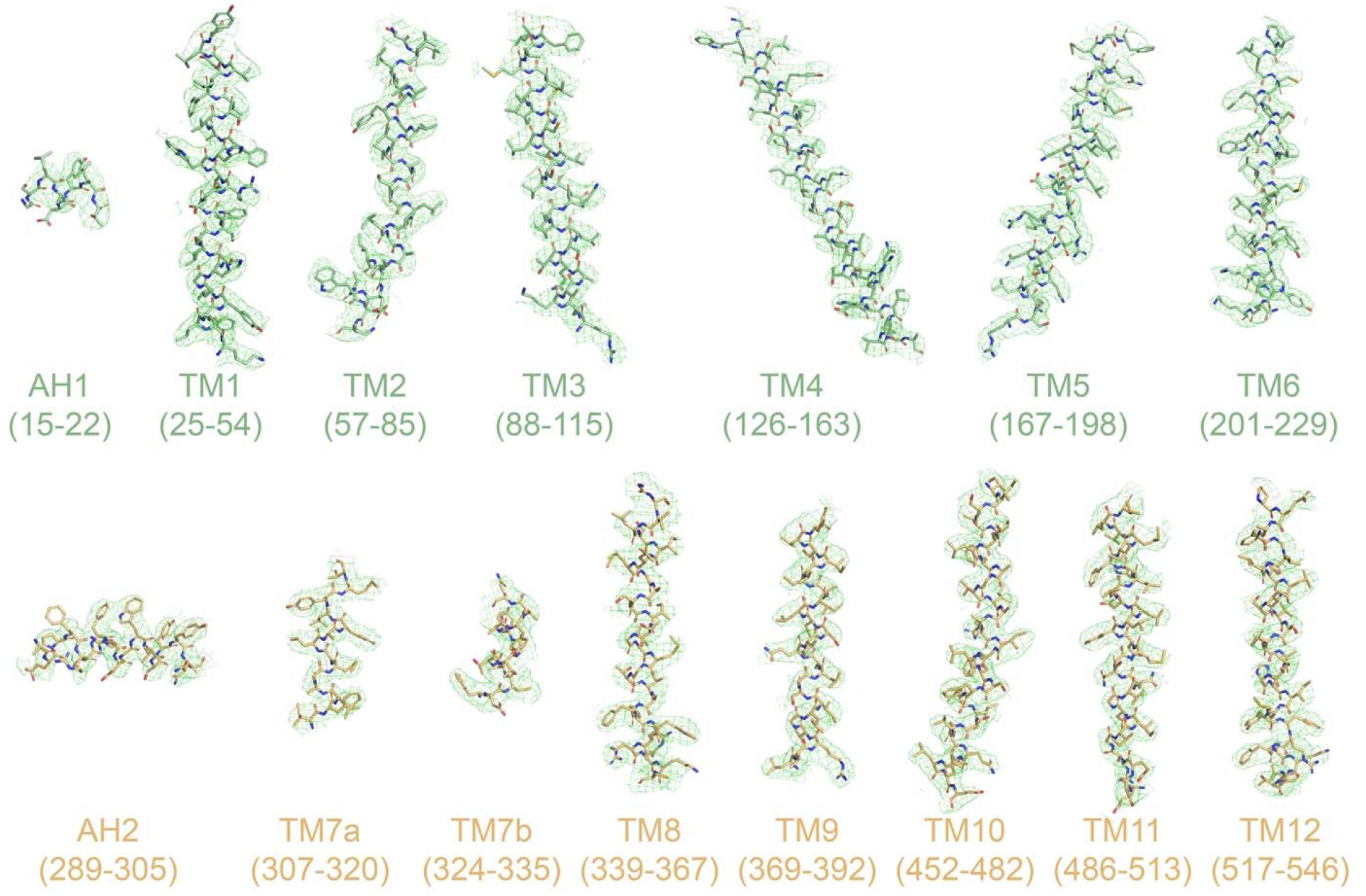
Cryo-EM densities of TM helices and amphipathic helices (AH). Densities for HsFpn-Ca^2+^ are shown as green mesh. Residues within the ranges indicated below are rendered in stick representation and colored in pale green for NTD or light orange for CTD.

**Figure 2—figure supplement 3.**
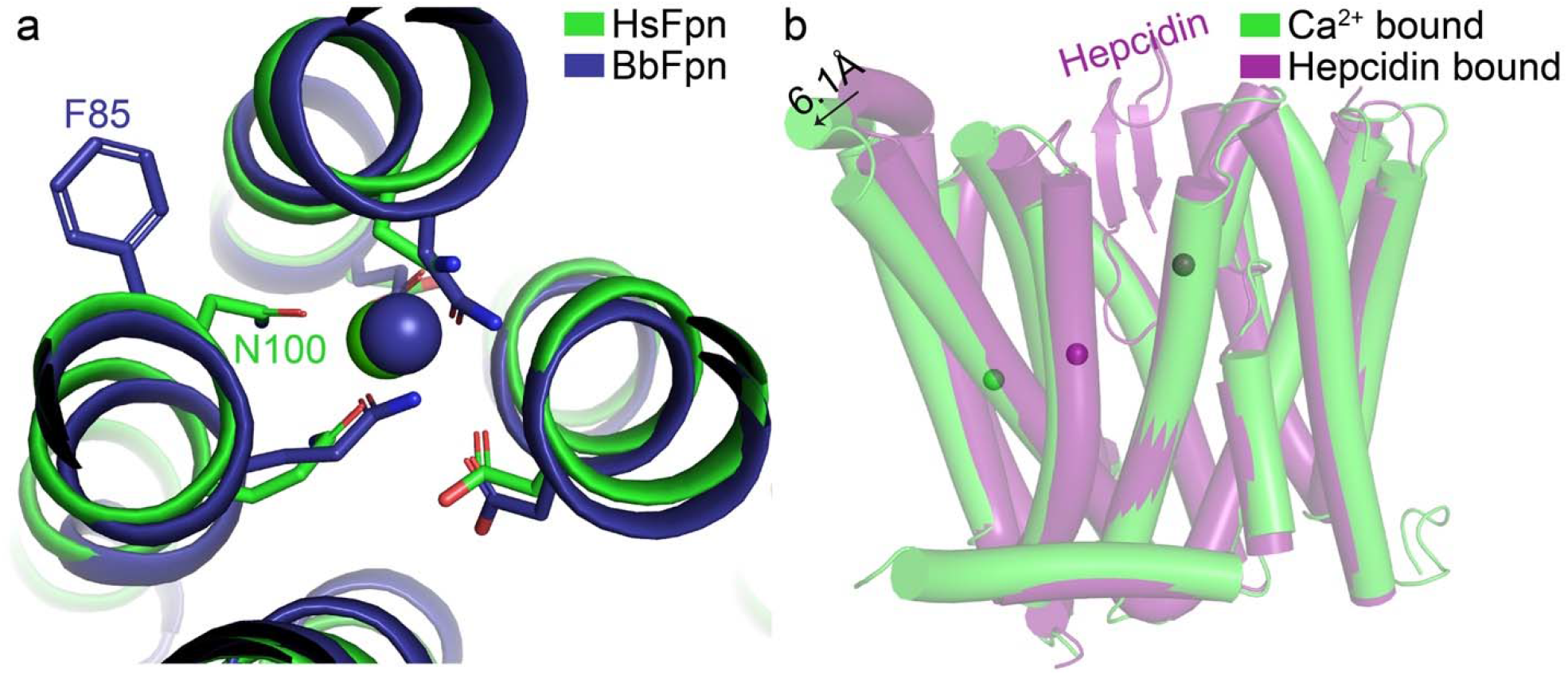
Structural comparisons of Ca^2+^-bound HsFpn. (**a**) Differences in Ca^2+^ coordination by HsFpn (green) and by BbFpn (deep blue, PDB ID 6BTX). (**b**) Comparison of Ca^2+^-bound (green) versus hepcidin-bound (deep purple, PDB ID 6WBV) structure of HsFpn shows further outward movements of NTD. Displacement of the loop between TM3 and TM4 is marked with an arrow and distance. Co^2+^ ions in the Fpn-hepcidin structure are shown as deep purple spheres.

**Figure 3—figure supplement 1.**
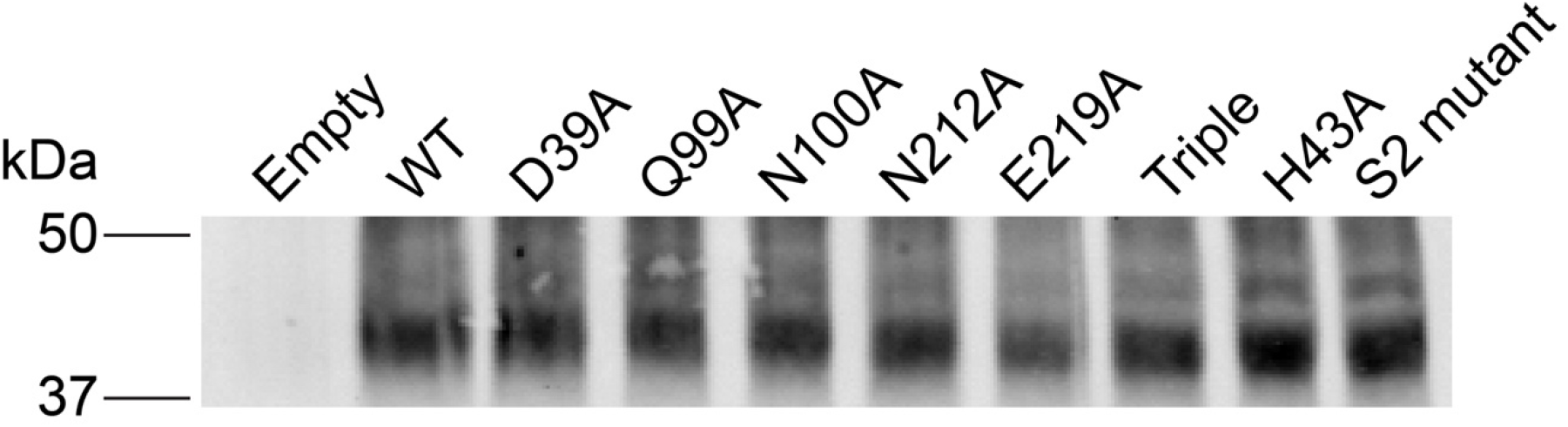
Expression of WT and mutant Fpn in HEK cells assessed by western blot.

**Figure 3—figure supplement 2.**
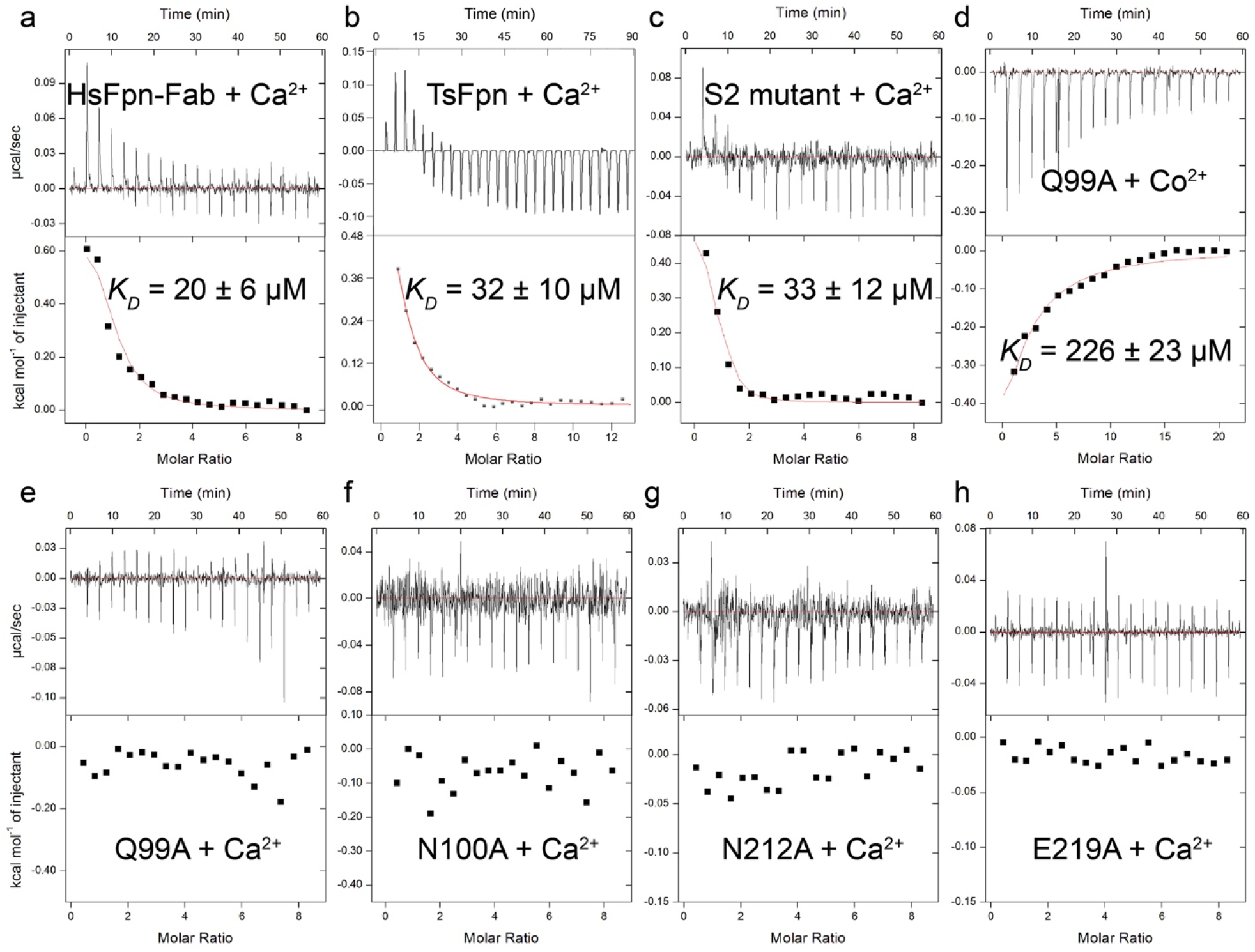
Binding of Ca^2+^ or Co^2+^ to WT and mutant Fpns measured by ITC. Binding of Ca^2+^ to: (**a**) HsFpn-11F9 complex; (**b**) TsFpn; (**c**) S2 mutant. (**d**) Binding of Co^2+^ to the alanine mutant of Q99 in the Ca^2+^ binding site. (**e**), (**f**), (**g**), and (**h**) Titration of Ca^2+^ into alanine mutants of Q99, N100, N212, and E219 in the Ca^2+^ binding site.

**Figure 4—figure supplement 1.**
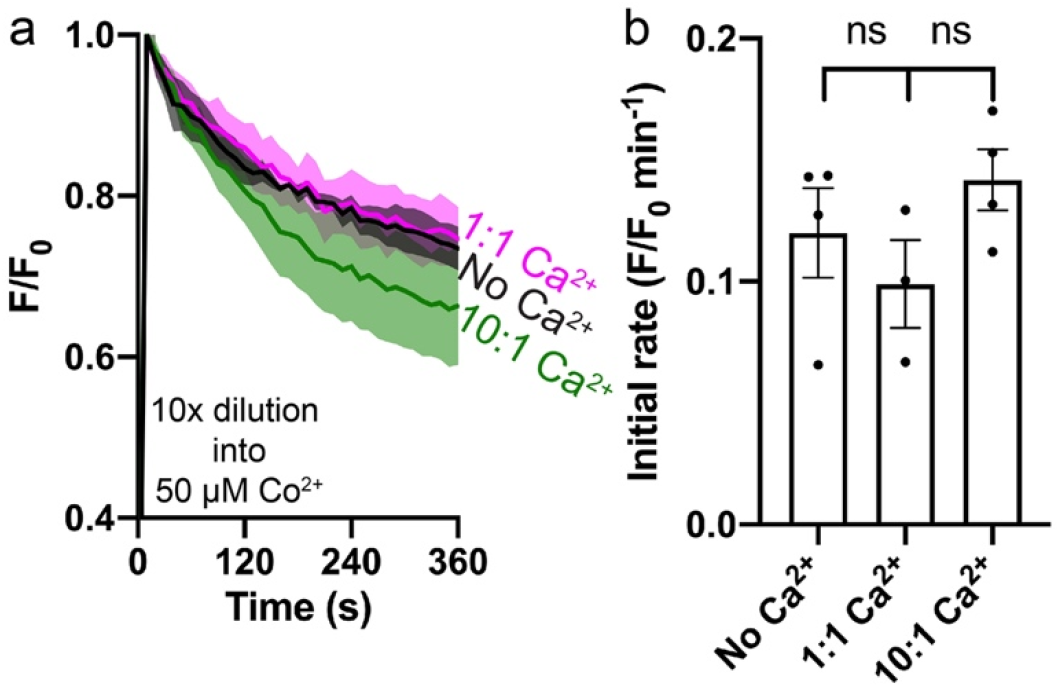
Co^2+^ transport by Fpn in the presence of Ca^2+^. (**a**) Co^2+^ influx in the presence of Ca^2+^ in proteoliposome. Liposome samples were diluted (10×) into outside buffers containing 50 µM Co^2+^. All fluorescence traces are subtracted from their corresponding no protein controls at the same conditions. (**b**) Comparison of initial rates of Co^2+^ transport in (**a**). One-way ANOVA was used for statistical analysis.

**Figure 4—figure supplement 2.**
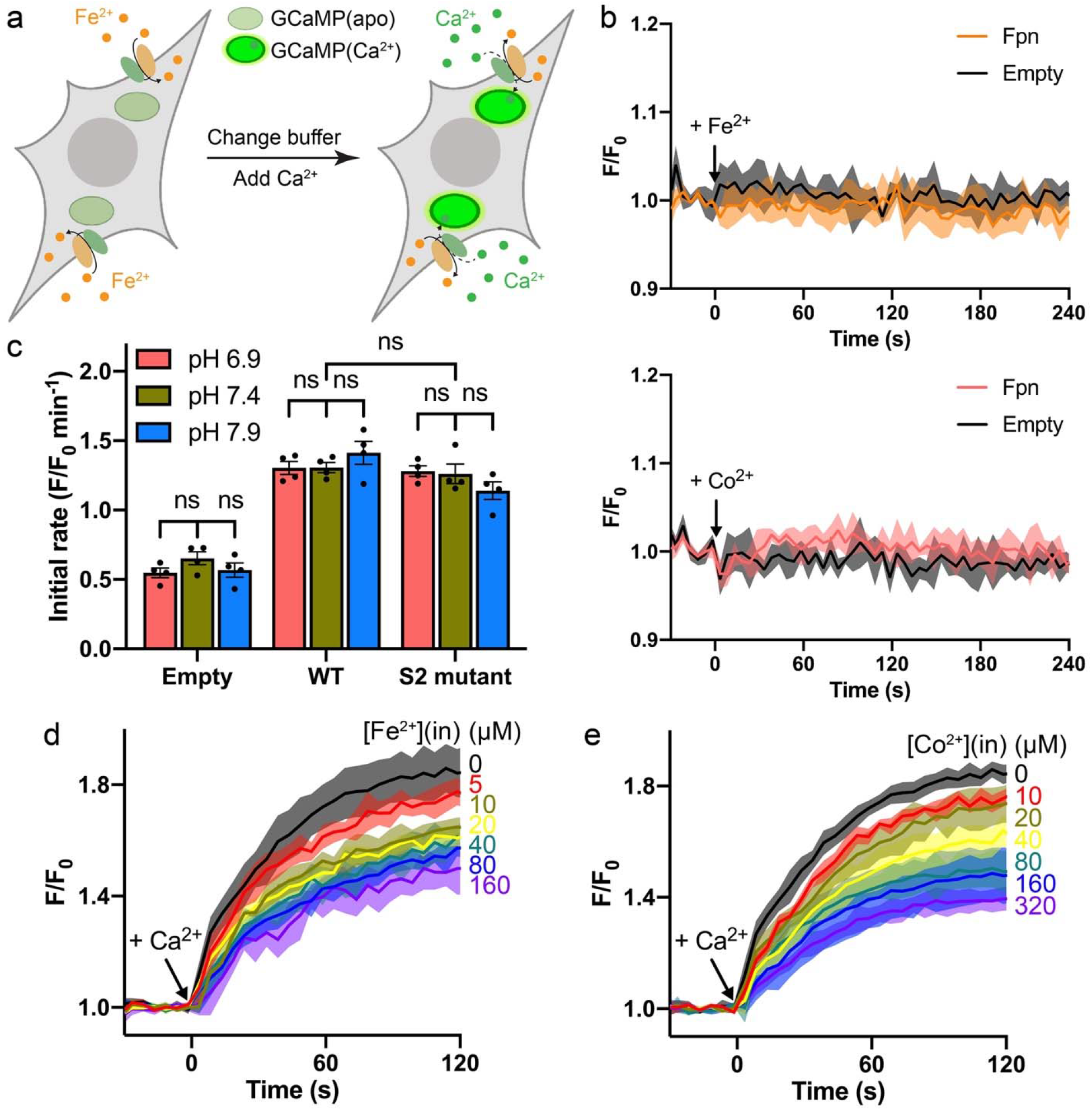
Ca^2+^ transport in the presence of Fe^2+^ or Co^2+^ in HEK cells. (**a**) Schematic of the Ca^2+^ transport assay in Fe^2+^-loaded HEK cells co-expressing Fpn and the Ca^2+^ sensor jGCaMP7s. (**b**) Transition metal ions do not interact with jGCaMP7s as indicated by unaltered fluorescence readings during loading of Fe^2+^ (top) or Co^2+^ (bottom) into HEK cells expressing Fpn (orange or pink traces) or transfected with empty vectors (gray traces). Black arrows indicate additions of metal ions. (**c**) Ca^2+^ transport at different extracellular pH in the presence of Co^2+^. Two-way ANOVA: among different pHs, p = 0.717; among empty, WT, and S2 mutant, p < 0.0001; interaction, p = 0.130. (**d**) and (**e**) Fpn-specific Ca^2+^ transport in the presence of Fe^2+^ or Co^2+^ loaded inside HEK cells. The concentrations of Fe^2+^ or Co^2+^ used during the loading step are indicated to the right of each trace. The Fe^2+^ stock was prepared with a 10-fold molar excess of sodium ascorbate. The raw traces of the empty control group were subtracted from the corresponding traces of the Fpn WT group. Normalized initial rates calculated from the datasets are shown in **Figure 4f–g**.

**Figure 4—figure supplement 3.**
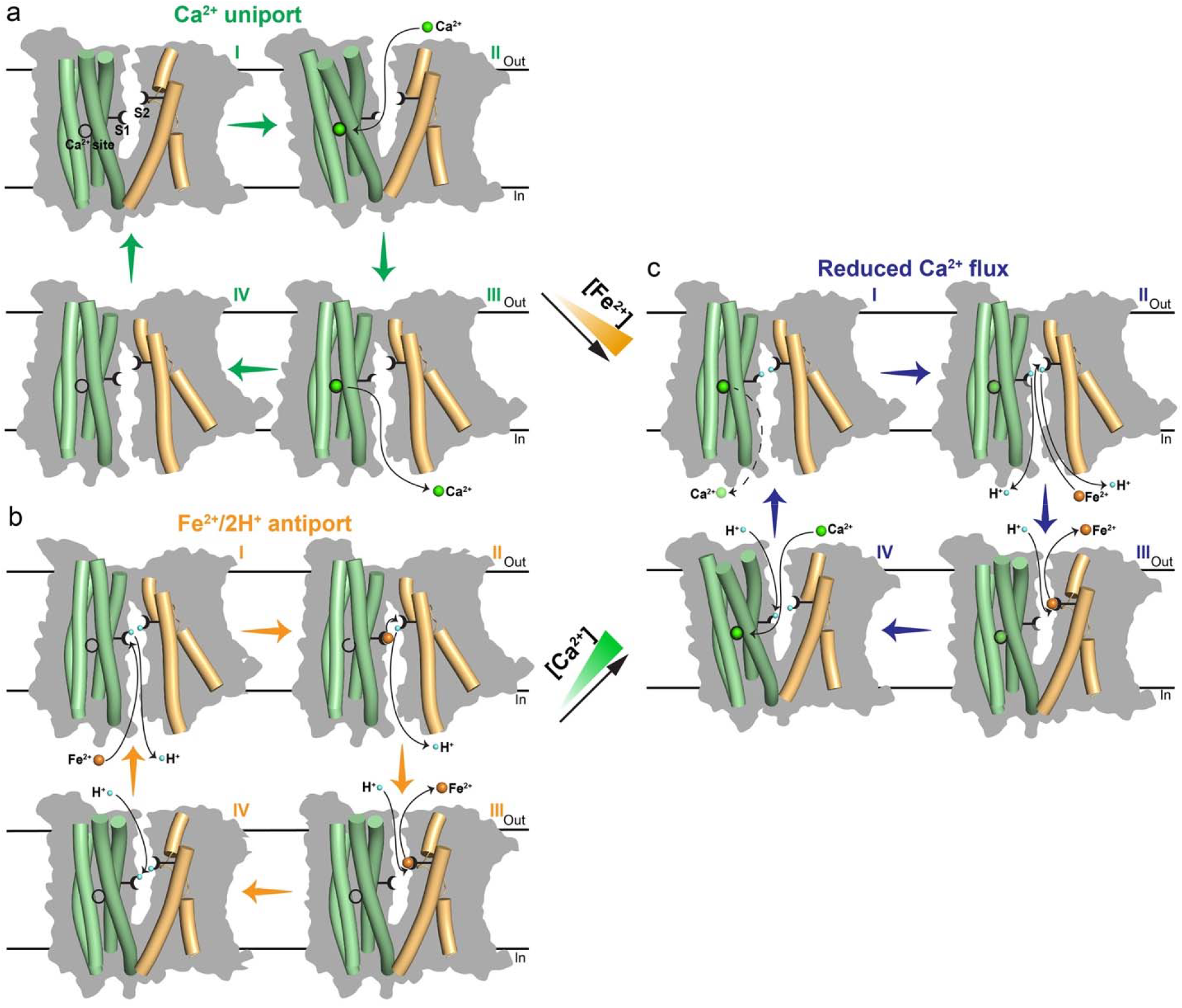
Proposed transport mechanisms of metal ions in Fpn. (**a**) Electrogenic Ca^2+^ uniport in the absence of Fe^2+^. (**b**) Electroneutral Fe^2+^/2H^+^ antiport in the absence of Ca^2+^. (**c**) Reduced Ca^2+^ flux in the presence of Fe^2+^. Higher concentration of Fe^2+^ (orange ramp) shows larger inhibition on Ca^2+^ transport. The inward-facing model is generated based on the bacterial homolog structure (PDB ID 6BTX). The four-helical bundle (pale green) in NTD, and TM7 and TM11 (light orange) in CTD are shown as cylinders and overlaid on silhouettes (grey) of the whole transporter. The Fe^2+^ S1 and S2 are indicated as black forks with two tines to represent two ligand residues in each site. The Ca^2+^ binding site is indicated as a black circle. When Fe^2+^ is bound to the S1, the Ca^2+^ binding site is drawn as an incomplete circle to indicate its partial disruption. Similarly, the S1 becomes a half fork when Ca^2+^ is bound. H^+^ (blue), Fe^2+^ (orange) and Ca^2+^ (green) are shown as spheres. The black dashed line and shaded green spheres of Ca^2+^ in **c** indicate reduced Ca^2+^ release and occupancy.

**Table 1.** Summary of cryo-EM data collection, processing, and refinement.

| Sample | HsFpn-Ca<sup>2+</sup>-11F9 |
| --- | --- |
| **Cryo-EM Data Collection** |  |
| Voltage (kV) | 300 |
| Magnification (×) | 81,000 |
| Pixel Size (Å) | 1.10 |
| Electron exposure (e<sup>-</sup>/Å<sup>2</sup>/frame) | 1.25 |
| Defocus range (μm) | [-2.5, -0.8] |
| Number of image stacks | 4,498 |
| Number of frames per stack | 40 |
| **Cryo-EM Data Processing** |  |
| Initial number of particles | 2,184,301 |
| Final number of particles | 437,959 |
| Symmetry imposed | C1 |
| Map resolution (Å) | 3.0 |
| Map resolution range (Å) | 2.4 – 3.6 |
| FSC threshold | 0.143 |
| **Model Refinement** |  |
| Number of amino acids | 875 |
| Total non-hydrogen atoms | 6,192 |
| Average B factor (Å<sup>2</sup>) | 171.4 |
| Bond length RMSD (Å) | 0.008 |
| Bond angle RMSD (°) | 0.939 |
| Ramachandran Plot |  |
| Favored (%) | 93.53 |
| Allowed (%) | 6.24 |
| Outliers (%) | 0.23 |
| Rotamer outliers (%) | 1.25 |
| MolProbity Score | 1.98 |

